# Allele frequency distribution of SNPs associated with levels of Vitamin D-binding protein and 25-hydroxyvitamin D

**DOI:** 10.1101/564229

**Authors:** Igor Nizamutdinov, Yaroslav Popov, Valery Ilinsky, Alexander Rakitko

## Abstract

Haplotypes defined by rs7041/rs4588 in GC gene modulate 25-hydroxyvitamin D (25(OH)D) and vitamin D-binding protein (DBP) levels. To investigate the distributions of GC polymorphisms, the rs7041 and rs4588 allele and haplotypes frequencies were analyzed in samples from different Eurasian regions. The GC1S haplotype associated with high level of serum 25(OH)D had the maximum frequency in European populations (except Finish population). Such frequency distributions may be a result of adaptation to low solar insolation rate. Besides, there was a strong trend of increasing GC1F haplotype frequency from Europe (10-15%) to Siberia and Easter Asia (40-45%).

## INTRODUCTION

Vitamin D deficiency is associated with negative health outcomes including cardiovascular disease (Skaaby, Thuesen, and Linneberg 2017), skeletal disorders (Wacker and Holick 2013) and type 2 diabetes mellitus (Harinarayan n.d.). After its synthesis in skin or absorption from dietary sources, vitamin D converts to 25-hydroxyvitamin D (25(OH)D). Serum total 25(OH)D is composed of three main metabolites: vitamin D-binding protein (DBP) bound form (85–90%), serum albumin bound form (10–15%) and free, unbound form (0.03%) (Bikle, Malmstroem, and Schwartz 2017) vitamin D status depends on the level of total 25(OH)D that is used to define vitamin D sufficiency.

DBP encoded by GC gene inhibits some actions of vitamin D, because the bound fraction may be unavailable to act on target cells (Safadi FF n.d.). Twin studies established that mature serum DBP concentration has a significant heritable component (62%) (Hunter D n.d.). Genetic variation in GC gene are shown to be associated with 25(OH)D levels (Wang et al. 2010). Recently, the racial difference was shown in DBP levels. Lower levels of DBP were shown in blacks compared to whites, although both groups had similar concentrations of estimated bioavailable 25(OH)D (Yousefzadeh, Shapses, and Wang 2014). However, the latest studies contradicted this racial difference and explained it by methodological faults (Nielson et al. 2016). It is still unclear whether DBP concentration relates to 25(OH)D level. Previous studies found no correlation between DBP and 25(OH)D levels (Bouillon R n.d.)(Winters SJ n.d.)(Blanton et al. 2011). The latest study revealed that high DBP level leads to low 25(OH)D level (Oleröd et al. 2017).

Two SNPs, rs7041 and rs4588, in GC gene form three haplotypes, namely GC1F, GC1S, and GC2, where GC1F = rs7041(A) and rs4588(G), GC1S = rs7041(C) and rs4588(G) and GC2 = rs7041(A) and rs4588(T). The combination of rs7041(C) and rs4588(T) does not exist in humans. The haplotypes were shown to have additive effects on DBP concentrations (Powe et al. 2013) and presumably on affinities between DBP and 25(OH)D (Arnaud and Constans 1993) (Chun et al. 2014). However, contrary results have been published that describe no association between SNPs and affinities between DBP and 25(OH)D (Boutin, Galbraith, and Arnaud 1989; R. Bouillon et al. 1984).

Blacks and Asians are more likely to carry GC1F haplotype, which is associated with low DBP levels and has the highest affinity for 25(OH)D. Whites are more likely to carry GC1S haplotype. The GC2 haplotype, which is associated with lower DBP levels and has a lower affinity for 25(OH)D, is frequently found in whites and rarely found in blacks (Chun RF n.d.; Powe CE n.d.).

Previous study found no differences in 25(OH)D values for children and adolescents, adults and elderly (Hilger et al. 2014), except for Asia/Pacific region where low 25(OH)D values found for Chinese children and adolescents. Also there were no significant sex-related differences in 25(OH)D before examining the data by region. Taking into account these data, we suggest that geographical region and GC polymorphisms are the main factors of population total 25(OH)D level difference.

We conducted a study to determine vitamin D–binding protein genotypes and haplotypes associated with vitamin D level in different populations.

## MATERIALS AND METHODS

### Data collection

The global rs7041 and rs4588 allele frequencies were drawn using GnomAD database (Lek et al. 2016). The rs7041 allele frequencies in different ethnic group were obtained from Alfred database (Rajeevan et al. 2012).

To investigate population differences in SNP frequencies in Eurasian populations, the genome data of Genotek Ltd. customers were analyzed. Totally, 4118 samples were studied with an average age of 33.3 ± 15.7. The group includes 46% female samples. All ethical requirements were covered by providing with informed consent of participants. Data used in our analysis were collected prior to February, 2018.

### Genotyping

Genotek Ltd. samples were drawn from the DNA biobank of Genotek Ltd., a direct-to-consumer genetics company. All samples were genotyped using customized Illumina InfiniumCoreExome 24 v1.1 beadchip with 550,000 standard plus 12,000 add-on SNPs and Illumina Global Screening Array v.1 and v.2.

### Statistical analysis

#### Imputing and phasing

Haplotypes were derived based on phasing using Beagle 5 (“Beagle 5.0” n.d.) with HRC panel (“The Haplotype Reference Consortium” n.d.) as the reference. Beagle 5 also was used for rs4588 imputation due to its absence on Illumina Global Screening Array.

#### Population inference

Ancestry was predicted using ADMIXTURE Version 1.3.0 in predictive mode (Alexander, Novembre, and Lange 2009). The samples from Behar et al. study (Behar et al. 2013) was used as original populations after exclusion of non-Ashkenazi jews. Each sample was assigned a population with maximal percent, but only if it was more than 50%.

## RESULTS

The global rs7041 and rs4588 allele frequencies are present in Table 1. The rs4588 (T) allele associated with increased levels of vitamin D–binding protein and decreased levels of total 25(OH)D in whites had the lowest frequency in African population. In whites, the lowest frequencies were observed in Finnish and Latino populations. The highest frequencies were shown for South Asian and Ashkenazi Jewish.

**Table 1.** Global rs4588 and rs7041 allele frequencies

| Geographic region | Sample size (n) |  | Allele Frequencies | Allele Frequencies |
| --- | --- | --- | --- | --- |
|  | rs4588 (G/T) | rs7041 (A/C) | rs4588-T | rs7041-A |
| European (non-Finnish) | 64540 | 64539 | 0.282 | 0.429 |
| European (Finnish) | 12556 | 9964 | 0.201 | 0.713 |
| Ashkenazi Jewish | 5185 | 5185 | 0.286 | 0.403 |
| East Asian | 9970 | 3607 | 0.271 | 0.446 |
| South Asian | 15287 | 12557 | 0.298 | 0.326 |
| Latino | 17663 | 17662 | 0.208 | 0.492 |
| African | 12474 | 12475 | 0.099 | 0.842 |
| Other | 3609 | 15292 | 0.262 | 0.449 |
| Total | 141284 | 141281 | 0.25 | 0.486 |

In contrast, rs7041 (A) allele associated with lower DBP levels and decreased levels of total 25(OH)D among blacks had the highest frequencies in Africans. In whites, the highest frequency was observed in Finnish population. Thus in African and Finnish populations, there were the highest frequencies of alleles associated with low DBP level.

To investigate allele frequency diversity within large geographical regions, the rs7041 allele frequencies were compared between different populations (Figure 1).

**Figure 1.**
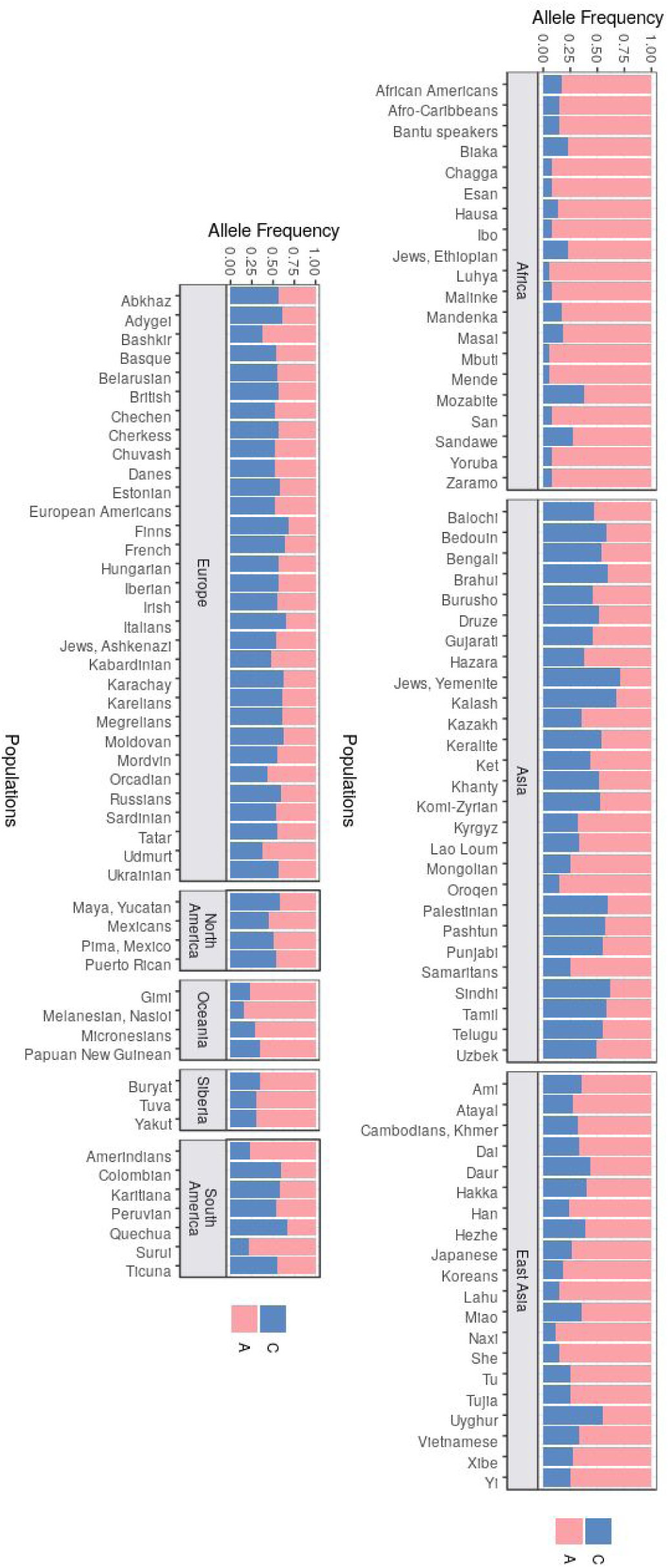
The rs7041 allele frequencies in different populations

Thereafter, the mean 25(OH)D level was obtained in various genotypes for different population (**STable 1**).

The allele and haplotype frequencies of rs7041 and rs4588 were estimated in a sample of 4118 Genotek Ltd customers. The obtained haplotype frequencies are present in **Table 2**. A maximum frequency was shown for haplotype GC1S associated with high level of serum 25(OH)D. The lowest frequency was observed for GC1F associated with lower level of serum 25(OH)D comparing with GC1S haplotype.

**Table 2.** Haplotype frequencies in Genotek biobank samples.

| Haplotype | Alleles | Frequency, % |
| --- | --- | --- |
| GC1S | rs7041(C)-rs4588(G) | 56.4 |
| GC2 | rs7041(A)-rs4588(T) | 29.9 |
| GC1F | rs7041(A)-rs4588(G) | 13.7 |
|  | rs7041(C)-rs4588(T) | 0 |

To investigate haplotypes distribution in Genotek Ltd samples divided in different populations, population origins were defined for 4118 DNA samples of Genotek Ltd biobank. Major part of samples had East Europe origin. Totally, nine clusters were defined in the sample set. The rs4588(T) and rs7041(A) allele frequencies are present on **SFigure 1** and **SFigure 2**, respectively.

The haplotype GC1F has low frequencies in all populations, except East Asian and Siberia (**SFigure 3**). Haplotype frequencies for GC1S are present on **Figure 2**. GC1S haplotype was the most frequent in all populations except East Asians and Siberia, where this haplotype had the lowest frequencies. Also there was a slight trend of increasing haplotype frequency from north to south.

**Figure 2.**
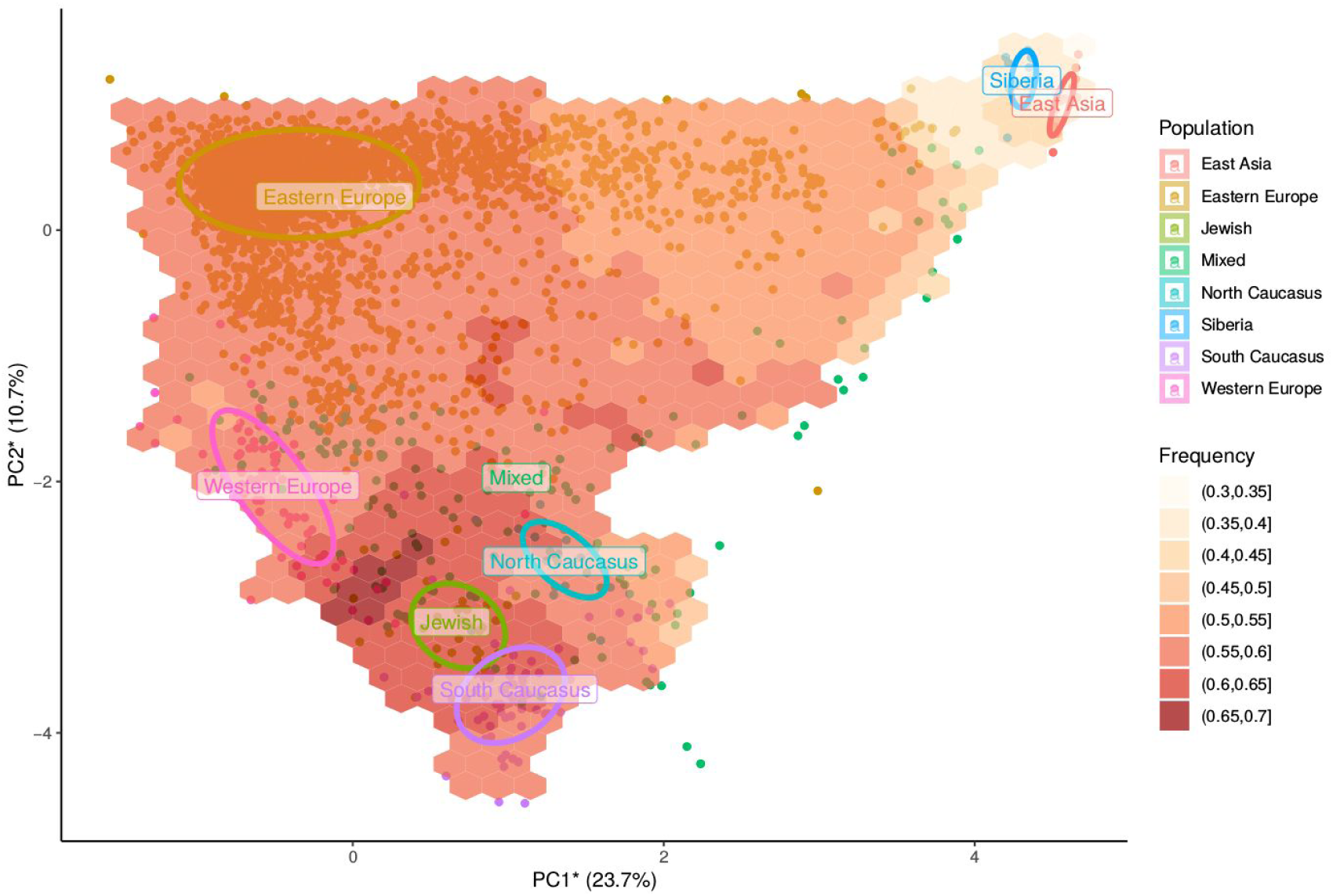
The GC1S haplotype frequencies in different Eurasian populations.

**Figure 3.**
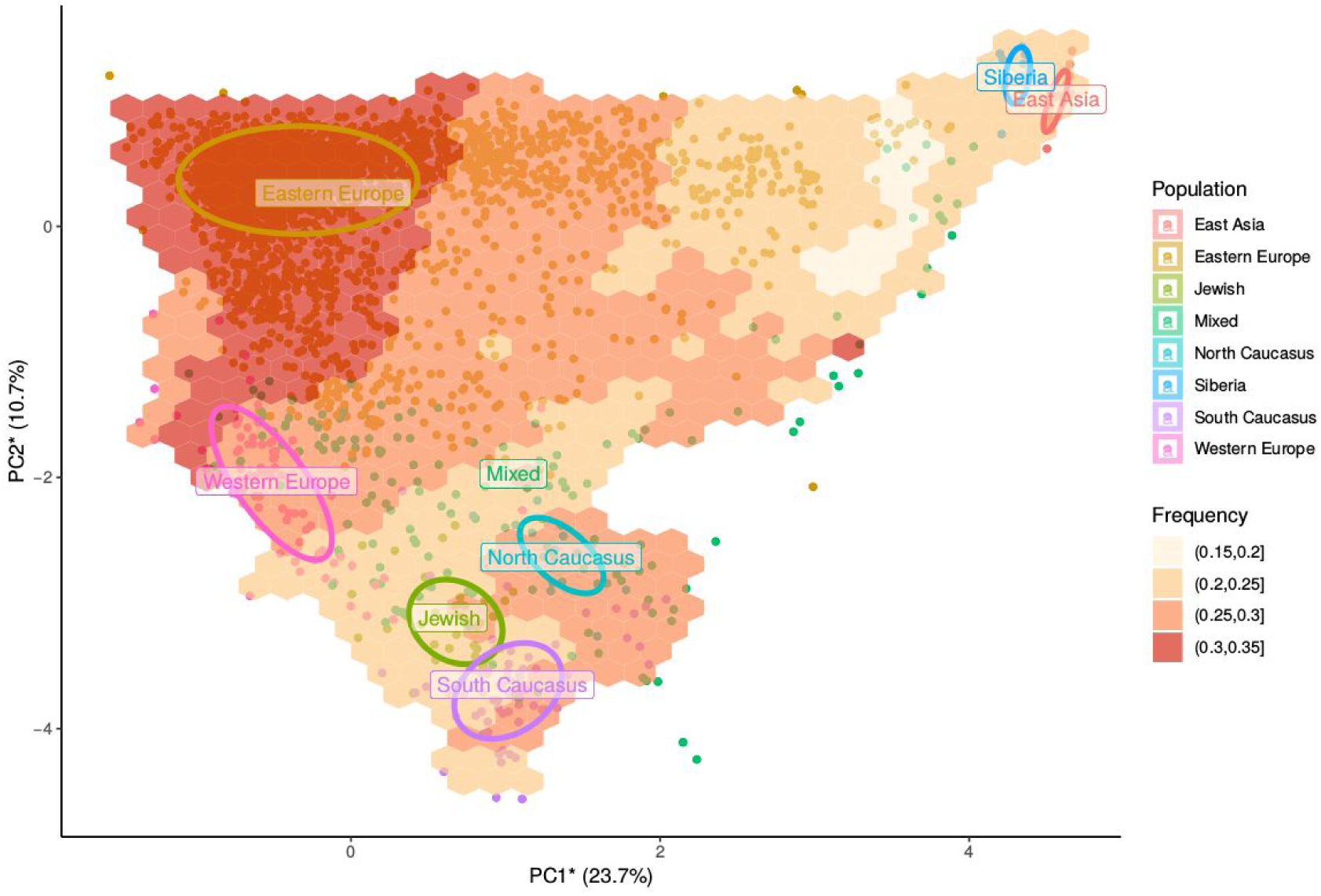
The GC2 haplotype frequencies in different Eurasian populations.

The highest GC2 frequency was shown for East Europe population. There were two trends of decreasing haplotype frequency, although the gradient was lower comparing with those of GC1F and GC1S. The GC2 frequency was lower for east populations (Siberia and East Asia). The second consists of decreasing haplotype frequency from north to south. There also was a difference in frequencies between Caucasus populations. North Caucasus population had a higher GC2 frequency comparing to South Caucasus population. Jewish population had the lowest GC2 frequency.

## DISCUSSION

Globally, there were maximum rs7041(A) allele frequency in African region. The Finnish population ranked second in rs7041(A) allele frequency. Recently, it was shown that blacks had the higher total 25(OH)D level comparing whites and Asians. At the same time, Finnish population had lower total 25(OH)D level comparing with most other European populations (Hilger et al. 2014). In Finns, the high frequency of rs7041(A) allele may be the consequences of a population bottlenecks resulting from consecutive founder effects (Chheda et al. 2017). In spite of rs7041 association with total 25(OH)D level in blacks, we suggest that genetic factors have minor influence on total 25(OH)D level in blacks.

Previous reviews (van Schoor and Lips 2011)(Lips 2007) reported a north-south gradient for 25(OH)D level in Europe, with Scandinavian countries showing generally higher values than European and Southern countries. We found no critical differences in rs7041(A) allele frequency in European populations except Finnish population. Probable these gradient is explained by non-genetic factors such as differences in skin pigmentation, diets rich in oily fish and low-fat milk, the common use of cod-liver oil and a higher degree of vitamin D supplementation in Scandinavian countries (Hagenau et al. 2009).

The GC1F haplotype had the lowest frequency in Genotek Ltd samples. However it was the most frequent in East Asia and Siberia population. Previously, it was shown that GC1F haplotype is the ancestral form of the gene (*Advances in Forensic Haemogenetics* 1994). This haplotype probably leads to the highest DBP affinity and lowest bioavailable 25(OH)D comparing with other haplotypes, what seems to be profitable in regions with high solar insolation rate. 25(OH)D bound form is inactive and perhaps could be a kind of storage of 25(OH)D overload (R. Bouillon and Van Baelen 1980).

The GC1S haplotype had the highest frequency in Genotek Ltd samples, except samples of East Asian and Siberia origin. Our results correspond to previous studies demonstrating dependence of GC1S frequency on the distance from the equator (Roger Bouillon 2017). GC1S variant occured at highest frequency at high latitudes. Vitamin D amount synthesized in skin is lower at high latitudes due to lower solar insolation rate. This haplotype is associated with lower DBP affinity comparing with GC1F and the highest levels of both total and bioavailable 25(OH)D comparing with other haplotypes (Yao et al. 2017), what seems to be adaptive factor to low vitamin D synthesis rate. Solar insolation rate is higher in Siberia and East Asia comparing with Europe, what seems to be a possible explanation of especial low GC1S level in these populations.

The frequency of GC2 haplotype was 29.9% in Genotek Ltd samples. The GC2 frequency variance was lower for different populations. This haplotype is associated with lowest DBP affinity comparing with other haplotypes and higher level of bioavailable 25(OH)D comparing with GC1F (Yao et al. 2017; Braithwaite et al. 2015), what could benefit in case of lower solar insolation rate.

Our study has several limitations. Not all populations living in Russia has been included in study. It is possible that small populations have different haplotype frequencies than large population groups. Research participants were drawn from customers of genetic company, what potentially could influence on sampling representativeness.

## Supplementary

**Table 1.** Mean 25(OH)Vitamin D Level (nmol/L) in the Various Genotypes obtained in different populations.

| Population | Number of patients | Sex | GC 1s/1s | GC 1f/1f | GC 2/2 |
| --- | --- | --- | --- | --- | --- |
| Norway (Jorde et al. 2015) | 11704 | both | 55.4 | 52.2 | 46.9 |
| Denmark (Fang et al. 2009) | 538 | both | 65.8 | 58.5 | 64.9 |
| French (Sinotte et al. 2009) | 727 | female | 67.5 | -- | 58.4 |
| Belgian (Janssens et al. 2010) | 150 | both | 65.75 | -- | 51.5 |
| Chinese (Zhang et al. 2013) | 2897 | both | 50.75 | -- | 42.25 |
| Finland (Pekkinen et al. 2014) | 160 | girls | -- | -- | 38.5 |
| Finland (Pekkinen et al. 2014) | 73 | boys | -- | -- | 45.3 |
| Jordanian (Lafi et al. 2015) | 200 | both | 90.25 | -- | 31 |
| **Mean** |  |  | **65.9** | **55.35** | **47.3** |

**Figure 1.**
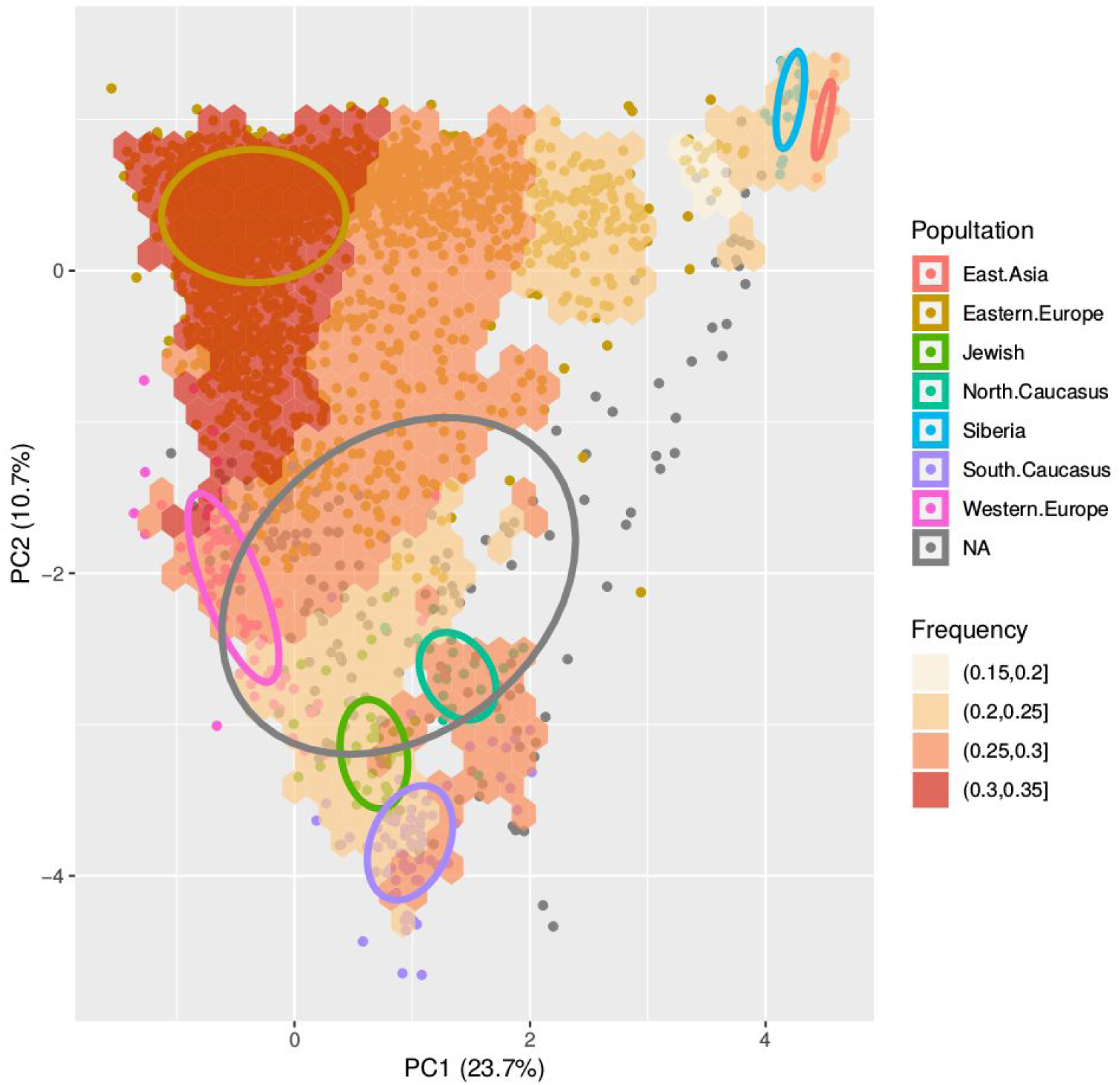
The rs4588(T) allele frequencies in different Eurasian populations. There were trends of decreasing allele frequency from west to east and from north to south. The highest T-allele frequency was observed in samples with east Europe origin. Lower T-allele frequency was shown for East Asia and Siberia populations. There was some difference of frequencies between Caucasus populations. North Caucasus population had a higher T-allele frequency comparing to South Caucasus population. Jewish population had low T-allele frequency.

**Figure 2.**
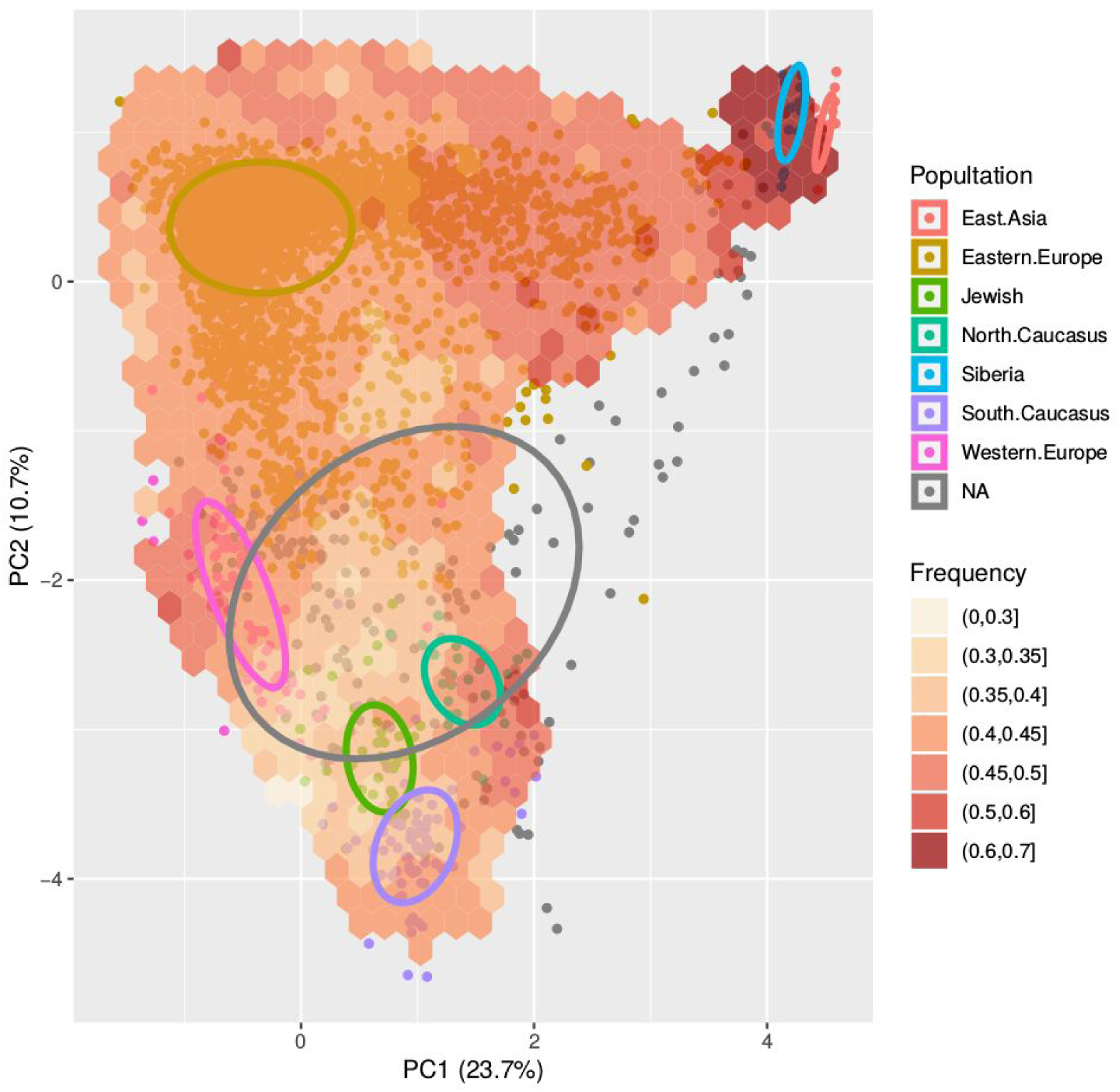
The rs7041(A) allele frequencies in different Eurasian populations. There was a trend of decreasing A-allele frequency from east to west and from north to south. Siberian and East Asian populations had the highest A-allele frequencies. There also was a difference of frequencies between Caucasus populations. North Caucasus population had a higher T-allele frequency comparing to South Caucasus population. Jewish population had the lowest T-allele frequency.

**Figure 3.**
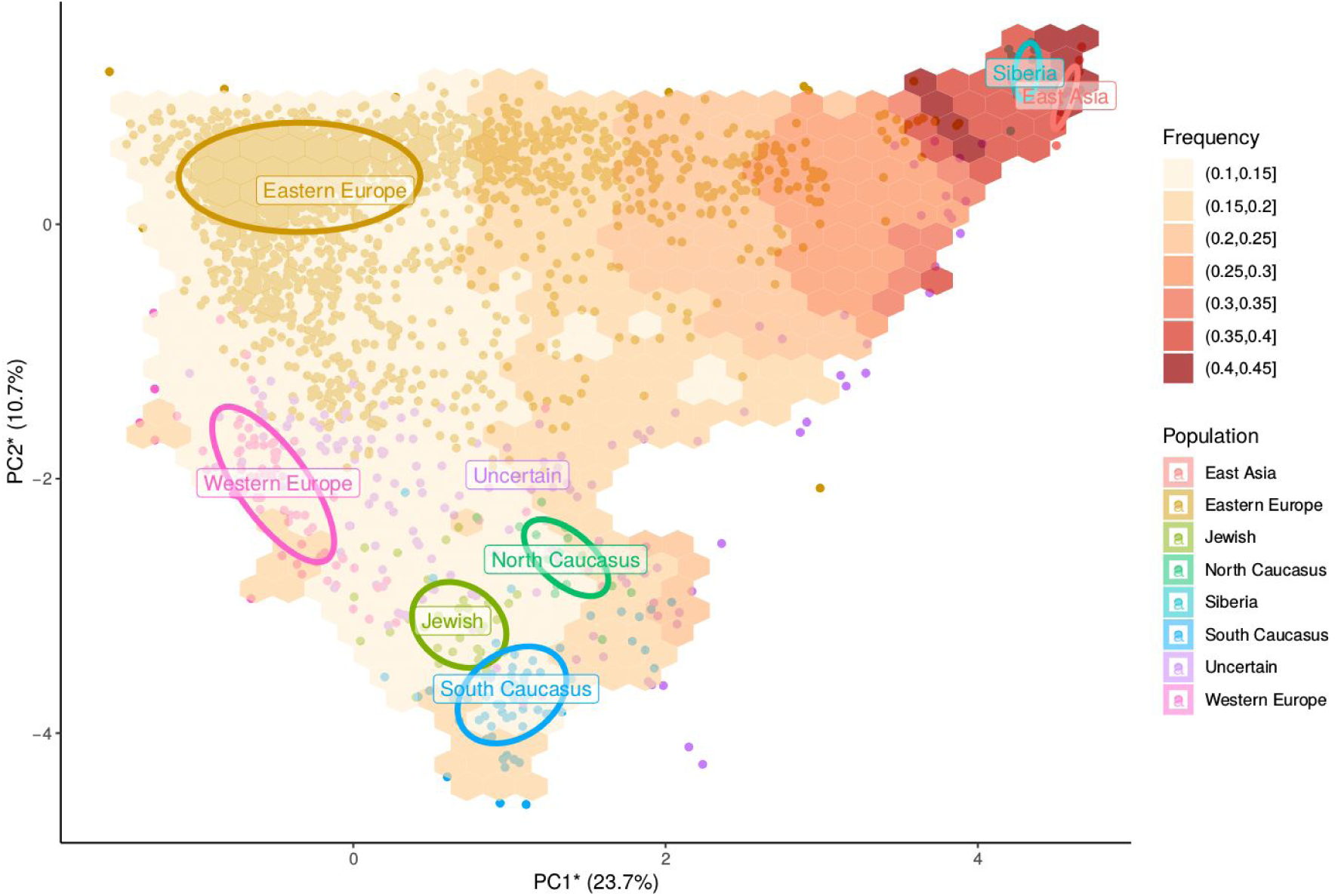
The GC1F haplotype frequencies in different Eurasian populations. European, Caucasus and Jewish populations had similar low frequencies. Siberian and East Asian populations had the highest A-allele frequencies.

